# Laminin N-terminus α31 expression during development is lethal and causes widespread tissue-specific defects in a transgenic mouse model

**DOI:** 10.1101/2020.07.26.221663

**Authors:** Conor J. Sugden, Valentina Iorio, Lee D. Troughton, Ke Liu, George Bou-Gharios, Kevin J. Hamill

## Abstract

Laminins are essential components of all basement membranes where they regulate an extensive array of tissue functions. Alternative splicing from the laminin α3 gene produces a non-laminin but netrin-like protein, Laminin N terminus α31 (LaNt α31). LaNt α31 is widely expressed in intact tissue and is upregulated in epithelial cancers and during wound healing. In vitro functional studies have shown that LaNt α31 can influence numerous aspects of epithelial cell behaviour *via* modifying matrix organisation, suggesting a new model of laminin auto-regulation. However, the function of this protein has not been established beyond the epithelium and it has never been studied in vivo. Here, a mouse transgenic line was generated using the ubiquitin C promoter to drive inducible expression of LaNt α31. When expression was induced at embryonic day 15.5, LaNt α31 transgenic animals were not viable at birth, exhibiting localised regions of erythema. Numerous striking defects were apparent histologically, including extra-vascular erythrocytes in multiple tissues, kidney epithelial detachment, tubular dilation, interstitial bleeding, and thickening of tubule basement membranes, disruption of the epidermal basal cell layer and of the hair follicle outer root sheath, and ~50% reduction of cell numbers in the liver associated with depletion of hematopoietic erythrocytic foci. These findings demonstrate that LaNt α31 can influence tissue morphogenesis during development. More broadly, these data provide the first in vivo evidence to support an emerging model of laminin self-regulation and provide a valuable model for onward investigation into this important area.

**Summary Statement:** Expression during development of Laminin N-terminus α31, a netrin-like laminin splice isoform, caused defects indicating basement membrane disruption in multiple tissues; providing the first in vivo evidence for laminin self-regulation.

## Introduction

Basement membranes (BMs) are specialised extracellular matrix (ECM) structures with essential and remarkably diverse roles in most cell and tissue behaviours; including regulating differentiation, cell adhesion and migration^1, 2^. BMs not only provide the mechanical attachment points that support sheets of cells to resist stresses but also influence signalling cascades via direct binding to cell surface receptors, through the sequestration and controlled release of growth factors, and by providing biomechanical cues, as reviewed in^3, 4^. BMs are also dynamic structures that are remodelled in terms of composition and structure throughout life, with the most striking changes occuring during development^5,6^. At the core of every BM are two networks of structural proteins; type IV collagens and laminins (LMs)^7^.

Each LM is an obligate αβγ heterotrimer formed from one of five α chains (*LAMA1-5*), three β chains (*LAMB1-3*) and three γ chains (*LAMC1-3*), with each chain displaying spatio-temporal distribution patterns, as reviewed in^8–11^. Assembly of LM networks and higher-order structures involves formation of a ternary node between the laminin N-terminal (LN) domains of an α, a β and a γ chain^12, 13^. These ternary αβγ nodes assemble in a two-step process involving an initial rapid formation of unstable βγ LN intermediate which is then stabilised through the incorporation of an α LN domain^14–17^. The biological importance of these LN-LN interactions is exemplified by a group of human syndromic disorders where missense mutations affecting the LN domains of the *LAMA2, LAMB2* or *LAMA5* genes give rise to muscular dystrophy in merosin-deficient muscular dystrophy, kidney and ocular developmental defects in Pierson syndrome, or defects in kidney, craniofacial and limb development respectively^18–22^. Although these disorders demonstrate that LM network assembly is essential for homeostasis of numerous tissues, not all LM chains contain an LN domain. Specifically, LMα4, which is expressed at high levels in the vasculature, and the LMα3a and LMγ2 chains, which are abundant in surface epithelium including the skin, have shortened amino termini which lack this key domain but yet still form functional BMs^8, 9, 23, 24^. This raises questions of whether LN domains are important in all tissue contexts or whether additional proteins may compensate for the inability of the LMs to form networks.

Alongside their main LM transcripts, the *LAMA3* and *LAMA5* genes produce short transcripts encoding proteins that are unable to trimerise into LMs but which contain LN domains^25^. At least one of these laminin N terminus proteins encodes a functional protein, LaNt α31, the structural features of which are an α LN domain followed by a short stretch of laminin-type epidermal growth factor-like (LE) domains and unique C-terminal region with no conserved domain architecture. In addition to the LaNt proteins, the laminin-superfamily includes netrin genes which encode proteins with either β or γ LN domains, stretches of LE repeats and unique C-terminal regions (as reviewed in^26^. Moreover, proteolytic processing of LMs are also released from LMα1^27^, LMβ1^28^, LMα31^29^. Each of these LN domain-containing proteins and cryptic fragments have cell surface receptor binding capabilities and can act as signalling molecules (reviewed in^30^. However, netrin-4, which evolved independently from the other netrins^31, 32^, also has LM-network disrupting capabilities^33,34^, and when overexpressed in vivo, caused increased lymphatic permeability^35^. Netrin-4 LN domain has greatest homology with LM β LN domains whereas LaNt α31 contains the LMα3b LN domain^36^; therefore, although LaNt α31 could act similarly to these proteins, it likely plays a different role depending on the LM context.

LaNt α31 is expressed in the basal layer of epithelia in the skin^25^, cornea^37^ and digestive tract, the ECM around terminal duct lobular units of the breast and alveolar air sacs in the lung, and is widely expressed by endothelial cells^38^. Increased expression is associated with breast ductal carcinoma and in vitro overexpression leads to a change in the mode of breast cancer cell invasion through LM-rich matrices^39^. LaNt α31 is also transiently upregulated during re-epithelialization ex vivo burn wounds and in stem cell activation assays^37^. In epidermal and corneal keratinocytes, knockdown or overexpression experiments revealed that modulating LaNt α31 levels leads to reduced migration rates and modifying cell-to-matrix adhesion^25, 40^. Consistent with a role in matrix assembly, increased expression LaNt α31 causes striking changes to LM332, including formation tight clusters beneath cells and increasing the proteolytic processing of LMα3 by matrix metalloproteinases^40^. Although these findings all support LaNt α31 as being a mediator of cell behaviour, it is as yet unknown what role it plays in complex in vivo tissue environments and in particualar in matrixes that are actively being remodeled.

Here, we present the first in vivo study of LaNt α31 overexpression in newly developed mouse models.

## Materials and methods

### Ethics

All procedures were licensed by the UK Home Office under the Animal (Specific Procedures) Act 1986, project license numbers (PPL) 70/9047 and 70/7288. All mice were housed and maintained within the University of Liverpool Biological Services Unit in specific pathogen-free conditions in accordance with UK Home Office guidelines. Food and water were available ad libitum.

### Antibodies

Rabbit monoclonal antibodies against the influenza hemagglutinin epitope (HA) (C29F4, Cell Signalling Technology, Danvers, MA) were used for immunoblotting at 67 ng ml^−1^. Goat polyclonal antibodies against DDDDK (equivalent to FLAG sequence, ab1257, Abcam, Cambridge, UK), rabbit polyclonal antibodies against 6X-His (ab137839, Abcam), and rabbit polyclonal antibodies against lamin A/C (4C11, Cell Signalling Technology) were used at 1 μg ml^−1^ for immunoblotting. Mouse monoclonal antibodies against LaNt α31^37^ were used at 0.225 μg ml^−1^ for immunoblotting. Rabbit polyclonal antibodies against mCherry (ab183628, Abcam) were used at 2.5 μg ml^−1^ for immunofluorescence. Alexa fluor 647 conjugated goat anti-rabbit IgG recombinant secondary antibodies, were obtained from Thermo Fisher Scientific (Waltham, MA, United States) and used at 2 μg ml^-1^ for indirect immunofluorescence microscopy.

### pUbC-LoxP-LaNtα31-T2A-tdTomato

A gBlock was synthesised (Integrated DNA Technologies, Coralville, IA) containing *Nde*I and *Nde*I restriction enzyme sites, T7 promoter binding site^41^, Kozak consensus sequence^42^, Igκ secretion signal (METDTLLLWVLLLWVPGSTGD)^43^, LaNt α31-encoding cDNA (amino acids 38-488)^25^, Flag (DYKDDDDK)^44^ and HA (YPYDVPDYA)^45^ tag sequences, T2A sequence (EGRGSLLTCGDVEENPGP)^46^, and *BamH*I. The gBlock DNA was inserted into pCSCMV:tdTomato (a gift from Gerhart Ryffel, Addgene plasmid #30530; http://n2t.net/addgene:30530; RRID:Addgene_30530) using *Nde*I and *BamH*I (New England Biolabs, Ipswich, MA), to produce pCS-LaNtø31-T2A-tdTomato. LaNtα31-T2A-tdTomato was then removed from this backbone using *Nhe*I and *EcoR*I and inserted into a vector containing the Ubiquitin C (UbC) promoter and a floxed stop cassette, all flanked by cHS4 insulator elements, producing pUbC-LoxP-LaNtα31-T2A-tdTomato.

### hK14-LaNt α31

Full length LaNt α31 cDNA was amplified by PCR and inserted into pSecTag vector (Thermo Fisher Scientific), introducing Igκ leader sequence 5’ of the LaNt α31 sequence, and Myc and 6x His tags 3’ of the LaNt α31 sequence. The complete Igκ-LaNt α31-Myc-His sequence was inserted into pGEM^®^-5Zf(+) vector (Promega, Madison, WI) using *Nhe*I and *Pme*I (New England Biolabs), producing pGEM^®^-5Zf(+)-LaNt α31. Separately, the sequence encoding human keratin 14 (hk14) promoter was amplified by PCR, using primers introducing *Mlu*I 5’ and *Nde*I, *Nsi*I 3’ of the sequence, and this was inserted into a bicistronic vector containing the mCherry seqeunce, producing phK14-mCherry. Finally, Igκ-LaNtα31-Myc-His was excised from pGEM^®^-5Zf(+)-LaNtα31 using *Nde*I and *Nsi*I (New England Biolabs) and inserted into phK14-mCherry, to produce phK14-LaNtα31-T2A-mCherry.

### Cloning procedures

Restriction digests were set up with 1 μg of plasmid DNA, 1 μg of PCR product, or 100 ng of gBlock DNA, 20 U of each enzyme and CutSmart buffer (50 mM Potassium Acetate, 20 mM Tris-acetate, 10 mM magnesium acetate, 100 μg ml^−1^ BSA (New England Biolabs) and incubated at 37°C for 1 h. Enzymatic activity was inactivated by 20 min incubation at 65°C. PCR or cloning products were separated using 1% (w/v) agarose gels (Thermo Fisher Scientific) dissolved in 1 x TAE electrophoresis buffer (40 mM Tris pH 7.6, 20 mM acetic acid, 1 mM EDTA) containing ethidium bromide, and visualised using a UV transilluminator ChemiDoc MP System (BioRad, Hercules, CA). DNA bands of the correct sizes were excised from the gel and purified using GenElute^™^ Gel Extraction Kit, following manufacturer’s protocol (Sigma Aldrich, St. Louis, Missouri, United States). Purified inserts were ligated into vectors at 3:1 molar ratios, either using Instant Sticky-end Ligase Master Mix (New England Biolabs) following manufacturers protocol, or using 400 U of T4 DNA ligase and 1X reaction buffer (50 mM Tris-HCl, 10 mM MgCl_2_ 1 mM ATP, 10 mM DTT, New England Biolabs) at 16°C overnight, followed by enzymatic inactivation at 65°C for 10 min. Ligated DNA was heat-shock transformed into One-Shot TOP10 chemically competent E. coli cells (Thermo Fisher Scientific) following manufacturer’s protocol, then plated onto LB plates containing the appropriate antibiotic (100 μg ml^−1^ ampicillin, 50 μg ml^−1^ kanamycin or 25 μg ml^−1^ chloramphenicol, Sigma Aldrich). Plasmid DNA was extracted from bacteria using GenElute^™^ Plasmid Miniprep Kit (Sigma Aldrich), following the manufacturer’s protocol. Plasmids were sequenced by DNASeq (University of Dundee, Dundee, UK).

### Cell Culture

KERA-308 murine epidermal keratinocyte cells^47^, were purchased from CLS (Cell Lines Service GmbH, Eppelheim, Germany) and maintained in high glucose (4.5 g L^−1^) Dulbecco’s Modified Eagle Medium (DMEM, Sigma Aldrich) supplemented with 10% foetal calf serum (LabTech, East Sussex, UK) and 2 mM L-glutamine (Sigma Aldrich). HEK293A cells were maintained in DMEM supplemented with 10% FCS and 4 mM L-glutamine.

### Cell Transfections

1 × 10□ KERA-308 or 4 × 10^5^ HEK293A cells were seeded in 6-well plates (Greiner-BioOne, Kremsmünster, Austria) 24 h prior to transfection. For KERA-308 cells, 2 μg of hK14-LaNto31-T2A-mCherry or LaNt-α31-pSec-Tag and 2 μl Lipofectamine 2000 (Thermo Fisher Scientific) were used. For HEK293A cells, either 1 μg pCAG-Cre:GFP and 2 μl Lipofectamine 2000, 2 μg of pUbC-LoxP-LaNtα31-T2A-tdTomato and 5 μl Lipofectamine 2000, or 2 μg of pUbC-LoxP-LaNtα31-T2A-tdTomato, 1 μg of pCAG-Cre:GFP and 7 μl Lipofectamine 2000 (Thermo Fisher Scientific), were mixed with 2 ml of Gibco™ Opti-MEM™ Reduced Serum Medium (Thermo Fisher Scientific) and incubated for 10 min at room temperature. The DNA-lipofectamine complex was added to the wells, and the media was replaced with DMEM high glucose after 6 h.

### Explant culture method

Hair was removed from mouse skin tissue using Veet hair removal cream (Reckitt Benckiser, Slough, UK) and the skin washed in Dulbecco’s Phosphate Buffered Saline (DPBS) containing 200 U ml^−1^ penicillin, 200 U ml^−1^ streptomycin, and 5 U ml^−1^ amphotericin B1 (all Sigma Aldrich). The skin was then dissected into 2-3 mm^2^ pieces using a surgical scalpel and 3 or 4 pieces placed per well of a 6-well dish (Greiner Bio-One, Kremsmünster, Austria) with the dermis in contact with the dish. 300 μl of DMEM supplemented with 20% FCS, 2 mM L-glutamine, 200 μg ml^−1^ penicillin, 200 μg ml^−1^ streptomycin, and 5 μg ml^−1^ fungizone (all Sigma Aldrich) was added to the wells. After 24 h, each well was topped up with 1 ml of media, and the media was replenished every 48 h thereafter.

### Transgenic Line establishment

Generation of transgenic mice were carried out based on the protocol described in^48^. C57Bl6CBAF1 females (Charles River Laboratories, Margate, Kent, UK) between 6-8 weeks were superovulated by intraperitoneal (IP) injections of 5 IU pregnant mare’s serum gonadotrophin (PMSG; in 100μl H_2_O), followed 46 h later by 5 IU of human chorionic gonadotropin (hCG, Sigma Aldrich). Treated females were mated with C57Bl6CBAF1 males overnight. Mated females were identified from the presence of copulation plugs, anaesthetised, and oviducts removed and dissected in M2 media (Millipore, Watford, UK). Day-1 oocytes (C57BL/6Jx CBA F1) were transferred into clean media by mouth pipetting. Cumulus cells were removed by hyaluronidase (300 μg ml^−1^, Merck, Darmstadt, Germany) treatment in M2 media with gentle shaking until detached from the egg surface. Oocytes were then rinsed and transferred to M16 media (Millipore, Speciality Media, EmbryoMax) ready for injection.

DNA was diluted to a final concentration of 2 ng μl^−1^ in embryo water (Sigma Aldrich) and filter-purified using Durapore-PVDF 0.22 μM centrifuge filters (Merck). Injection pipettes were used to pierce the outer layers of the oocyte and to inject DNA. DNA was injected into the pronuclei of the oocyte. Undamaged eggs were transferred to clean M16 media and incubated at 37°C until transferred into pseudopregnant CD1 females on the same day. Meanwhile, pseudopregnant females were obtained by mating vasectomised CD1 males overnight. Copulation plugs were checked and females were used 1 day post-coitum. Females were anaesthetised by inhalation of isoflurane (Sigma Aldrich). 30 injected oocytes were transferred to plugged pseudopregnant female oviducts through the infundibulum.

In generating the pUbC-LoxP-LaNtα31-T2A-tdTomato line, 460 mouse zygotes were injected over four sessions. 87% of these zygotes survived and were transferred into 11 recipient CD1 mothers. From these mothers, 42 pups were born. Of the 10 F0 mice that gave a positive genotype result, four passed on the transgene to the F1 generation. Mice that did not pass on the transgene to the F1 generation were culled, the four F0 mice were mated to expand colonies for cryopreservation, and one line was continued for investigation.

For K14-LaNtα31 transgenic mice, 140 embryos were transferred into five recipient CD1 mothers. Three small litters were born, totalling seven pups. two pups possessed the transgene, and these were mated to generate F1 mice.

R26CreERT2 (Jax Lab 008463)^49^ mice were purchased from The Jackson Laboratory (Bar Harbor, Maine, United States).

### In Vivo Transgene Induction

Tamoxifen (Sigma Aldrich) was dissolved in corn oil (Sigma Aldrich) and administered via IP at a concentrations of 25 mg kg^−1^ or 75 mg kg^−1^. Progesterone (Sigma Aldrich) was dissolved in corn oil (Sigma Aldrich) and was co-administered alongside tamoxifen at a dose of exactly half of the corresponding tamoxifen dose (12.5 mg kg^−1^ or 25 mg kg^−1^).

### DNA Extraction

Four weeks after birth, ear notches were collected from mouse pups and digested in 100 μl lysis buffer (50 mM Tris-HCl pH 8.0, 0.1 M NaCl, 1% SDS, 20 mM EDTA) and 10 μl of proteinase K (10 mg ml^−1^, all Sigma Aldrich) overnight at 55°C. The following day, samples were cooled, spun at 13,000 rpm for 3 min and the supernatant transferred to clean 1.5 ml tubes (Eppendorf, Hamburg, Germany). An equal volume of isopropanol (Sigma Aldrich) was added, gently inverted and span at 13,000 rpm, and supernatant discarded. Pellets were washed with 500 μl of 70% EtOH (Sigma Aldrich), then air-dried for 10 min, and resuspended in 50 μl ddH_2_0.

### PCR

50 ng of genomic DNA was mixed with 12.5 μl of REDtaq ReadyMix PCR Reaction Mix (20 mM Tris-HCl pH 8.3, 100 mM KCl, 3 mM MgCl2, 0.002% gelatin, 0.4 mM dNTP mix, 0.06 unit/ml of Taq DNA Polymerase, Sigma Aldrich) and 0.5 μM of each primer; ddH20 was added to make the reaction mixture up to 25 μl. Primer pairs for genotyping were as follows: LaNt α31 to tdTomato Forward 5’ – ATCTATGCTGGTGGAGGGGT – 3’, Reverse 5’ – TCTTTGATGACCTCCTCGCC – 3’; Cre Forward 5’ – GCATTACCGGTCGATGCAACGAGTGATGAG – 3’, Reverse 5’ – GAGTGAACGAACCTGGTCGAAATCAGTGCG – 3’; Recombination Forward 5’ – TCCGCTAAATTCTGGCCGTT – 3’, Reverse 5’ – GTGCTTTCCTGGGGTCTTCA – 3’(all from Integrated DNA Technologies). Cycle conditions were as follows: Genotyping – 1 cycle of 95°C for 5 min, 35 cycles of 95°C for 15 s; 56°C for 30 s; 72°C for 40 s, followed by a final cycle of 72°C for 5 min. For checking recombination: 1 cycle of 95°C for 5 min, 35 cycles of 95°C for 15 s; 60°C for 30 s; 72°C for 90 s, followed by a final cycle of 72°C for 7 min. PCR products were separated by gel electrophoresis and imaged using a BioRad Gel Doc XR+ System.

### SDS-PAGE and western immunoblotting

Cells were homogenized by scraping into 90 μL Urea/SDS buffer: 10 mM Tris-HCl pH 6.8, 6.7 M urea, 1% w/v SDS, 10% v/v glycerol and 7.4 μM bromophenol blue, containing 50 μM phenylmethysulfonyl fluoride (PMSF) and 50 μM N-methylmaleimide (all Sigma Aldrich). Lysates were sonicated and 10% v/v β-mercaptoethanol (Sigma Aldrich) added. Proteins were separated by sodium dodecyl sulfate-polyacrylamide gel electrophoresis (SDS-PAGE) using 10% polyacrylamide gels; 1.5 M Tris, 0.4% w/v SDS, 10% acrylamide/ bis-acrylamide (all Sigma Aldrich), electrophoresis buffer; 25 mM tris-HCl, 190 mM glycine, 0.1% w/v SDS, pH 8.5 (all Sigma Aldrich). Proteins were transferred to a nitrocellulose membrane using the TurboBlot™ system (BioRad) and blocked at room temperature in Odyssey^®^ TBS-Blocking Buffer (Li-Cor BioSciences, Lincoln, NE, United States) for 1 h. The membranes were probed overnight at 4°C diluted in blocking buffer, washed 3 × 5 min in PBS with 0.1% Tween (both Sigma Aldrich) and probed for 1 h at room temperature in the dark with IRDye^®^ conjugated secondary Abs against goat IgG (800CW) and rabbit IgG (680CW), raised in goat or donkey (LiCor BioSciences), diluted in Odyssey^®^ TBS-Blocking Buffer at 0.05 μg ml^−1^. Membranes were then washed for 3 × 5 min in PBS with 0.1% Tween, rinsed with ddH_2_O and imaged using the Odyssey^®^ CLX 9120 infrared imaging system (LiCor BioSciences). Image Studio Light v.5.2 was used to process scanned membranes.

### Tissue processing

For cryosections, P0 pups were culled by cervical dislocation, and fixed in 4% paraformaldehyde (Merck) overnight at 4°C. Samples were cryoprotected in 30% sucrose/PBS solutions then in 30% sucrose/PBS:O.C.T (1:1) solutions (Tissue-Tek, Sakura Finetek Europe, Alphen aan den Rijn, The Netherlands), each overnight at 4°C. Samples were embedded in OCT compound and transferred on dry ice. Embedded samples were sectioned at 8 μm using a Leica CM1850 cryostat (Leica, Wokingham, UK). For paraffin sections, Tissues were fixed in 10% neutral buffered formalin (Leica,) for 24 h, then processed through graded ethanol and xylene before being embedded in paraffin wax. 5 μm sections were cut using a rotary microtome RM2235 (Leica), adhered to microscope slides, then dried overnight at 37°C. Sections were dewaxed and rehydrated with xylene followed by a series of decreasing ethanol concentrations. Antigen retrieval was performed by microwaving sections in preheated 0.01 M citrate buffer pH 6 (Sigma Aldrich) for 5 min.

### Hematoxylin and Eosin Staining

Sections were dewaxed and rehydrated with xylene followed by a series of decreasing ethanol concentrations. Sections were then stained in Harris hematoxylin solution (Leica) for 5 min, H_2_O for 1 min, acid alcohol (Leica) for 5 s, H_2_O for 5 min, aqueous eosin (Leica) for 3 min, H_2_O for 15 s, followed by dehydration through graded ethanol and xylene. Slides were coverslipped with DPX mounting media (Sigma Aldrich).

### Immunohistochemistry

Slides were incubated in ice-cold acetone for 10 min, then transferred into PBS for 10 minblocking, then blocked in PBS containing 10% normal goat serum (NGS) at room temperature for 1 h. Next, samples were probed with the primary antibodies diluted in PBS-Tween (0.05%) with 5% NGS at 4°C overnight. Samples were then washed for 3 × 5 min in PBS-Tween (0.05%), before being probed with secondary antibodies diluted in PBS-Tween (0.05%) with 5% NGS at room temperature for 1 h. Samples were washed for 3 x 5 min in PBS-Tween (0.05%). Slides were mounted with VECTASHIELD^®^ Antifade Mounting Medium with DAPI (VECTASHIELD^®^, Burlingame, CA).

### Image Acquisition

H&E images were acquired using a Zeiss Axio Scan.Z1 equipped with an Axiocam colour CCD camera using ZEN Blue software (all from Zeiss, Oberkochen, Germany). Live cell images were acquired using a Nikon Eclipse Ti-E microscope (Nikon, Tokyo, Japan). Immunofluorescence images of tissues were acquired using a Zeiss LSM 800 confocal microscope (Zeiss).

### Image Analysis

Images were processed using either Zen 2.6 (blue edition) (Zeiss) or ImageJ (National Institutes of Health, Bethesda, MD, United States)^50^. Stardist plugin^51^ was used for segmentation of nuclei from H&E images. Images were thresholded manually to remove areas containing no tissue in the images.

## Results

### Inducible LaNt α31 construct validation

To investigate the consequences of LaNt α31 overexpression in vivo, we generated an inducible system for conditional LaNt α31 transgene expression (Fig. 1A). An expression construct was created containing the ubiquitin C promoter driving expression of the human LaNt α31 cDNA with the native secretion signal replaced by mouse immunoglobulin κ leader sequence to maximise secretion, and with sequences for Flag and HA epitope tags added to the C-terminus of the LaNt α31 coding region. A T2A element was included to enable expression of tdTomato from the same transgene but not directly fused to LaNt α31^46^. A floxed stop-cassette was inserted between the promoter and the start of the construct to prevent transgene expression until Cre-mediated removal of this cassette. The entire construct was flanked with the cHS4 β-globin insulator to protect against chromatin-mediated gene silencing^52^ (Fig. 1A). Restriction enzyme digests and plasmid sequencing confirmed the assembled pUbC-LoxP-LaNtα31-T2A-tdTomato plasmid.

**Figure 1.**
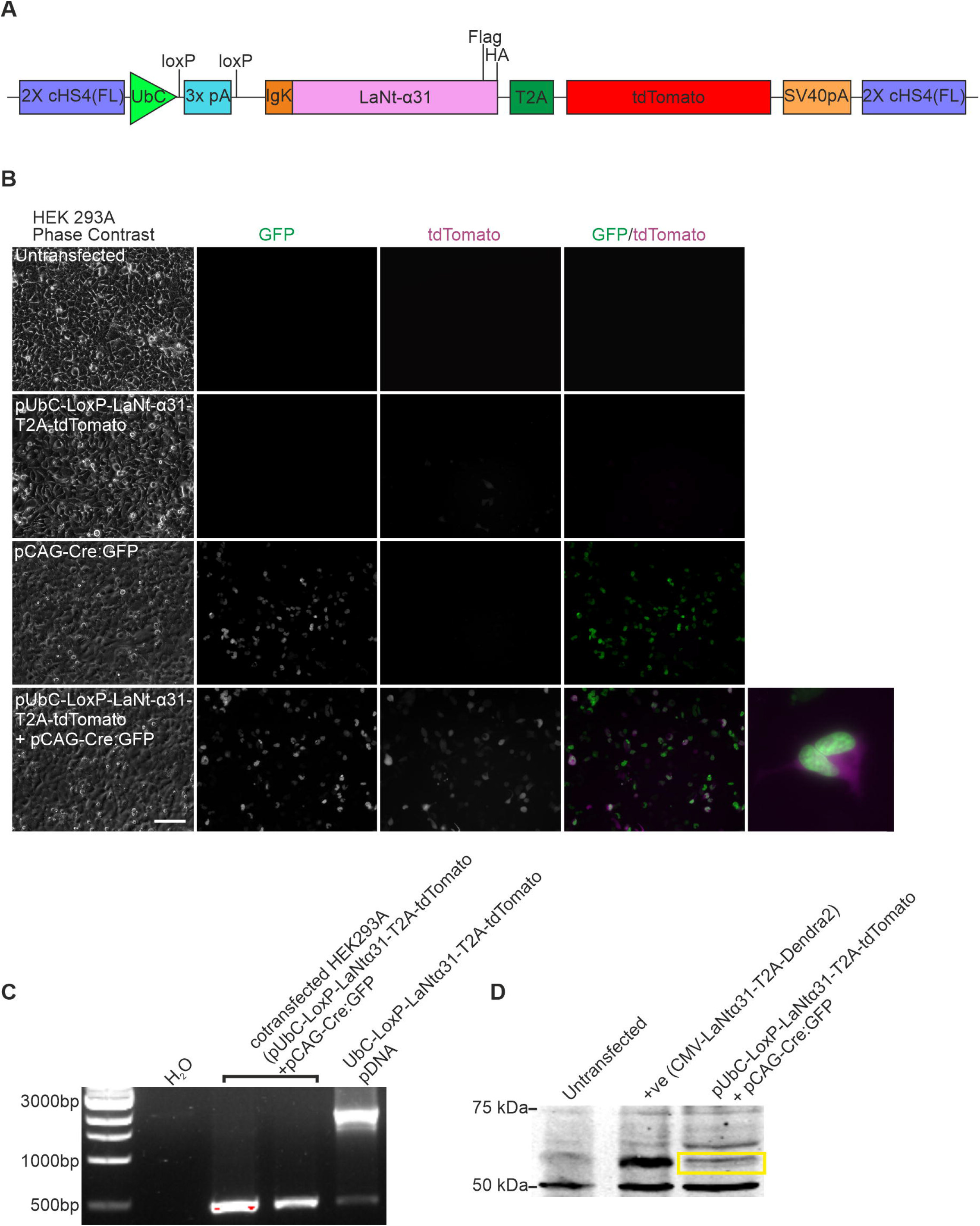
Validation of UbCLaNt Cre-inducible construct in vitro. A) Diagram of the pUbC-LoxP-LaNt-α31-T2A-tdTomato construct. B) HEK 293A cells were transfected with pUbC-LoxP-LaNt-α31-T2A-tdTomato, pCAG-Cre:GFP, or pUbC-LoxP-LaNt-α31-T2A-tdTomato and pCAG-Cre:GFP and imaged 48 h after transfection. Scale bar 100 μm C) PCR was performed using primers flanking the stop cassette on DNA extracted from HEK293A cells co-transfected with pUbC-LoxP-LaNt-α31-T2A-tdTomato and pCAG-Cre:GFP. D) Western blot of lysates from HEK293 cells either untransfected or transfected with CMV-LaNt-α31-T2A-Dendra2 (positive control), or pUbC-LoxP-LaNt-α31-T2A-tdTomato and pCAG-Cre:GFP then probed with anti-flag antibodies

To confirm the construct expressed only following exposure to Cre recombinase, the pUbC-LoxP-LaNtα31-T2A-tdTomato was co-transfected alongside pCAG-Cre:GFP, encoding GFP-tagged Cre recombinase, into HEK293A cells. tdTomato signal was observed only in cells transfected with both plasmids (Fig. 1B). PCR using primers flanking the STOP cassette also confirmed that the cassette was removed only in cells transfected with both plasmids (Fig. 1C). Western blotting using polyclonal anti-Flag antibodies confirmed expression of the predicted ~ 57 kDa band in co-transfected cell lysates (Fig. 1D), this also confirmed that the T2A element was cleaved in the final product releasing the tdTomato tag. Together, these results demonstrate that the pUbC-LoxP-LaNtα31-T2A-tdTomato plasmid allows for the Cre-inducible expression of LaNt α31 and tdTomato.

### Generation and validation of a novel LaNt α31 overexpressing mouse line

The pUbC-LoxP-LaNtα31-T2A-tdTomato construct was linearised, and transgenic F0 mice generated by pronuclear microinjection into oocytes. To confirm transgene expression, F0 mice were mated with WT (C57BL/6J) mice, embryos were collected at E11.5, and mEFs were isolated from the embryos. Presence of the UbC-LoxP-LaNto31-T2A-tdTomato transgene (hereafter UbCLaNt) was confirmed by PCR (Fig. 2A). mEFs were transduced with an adenovirus encoding codon-optimised Cre recombinase (ad-CMV-iCre). Analysis by immunoblotting with anti-HA-antibodies (Fig. 2B) revealed a ~57 KDa band and fluorescence microscopy confirmed tdTomato expression (Fig. 2C) in samples containing both the UbCLaNt transgene and the ad-CMV-iCre, but not in cells with either plasmid individually.

**Figure 2.**
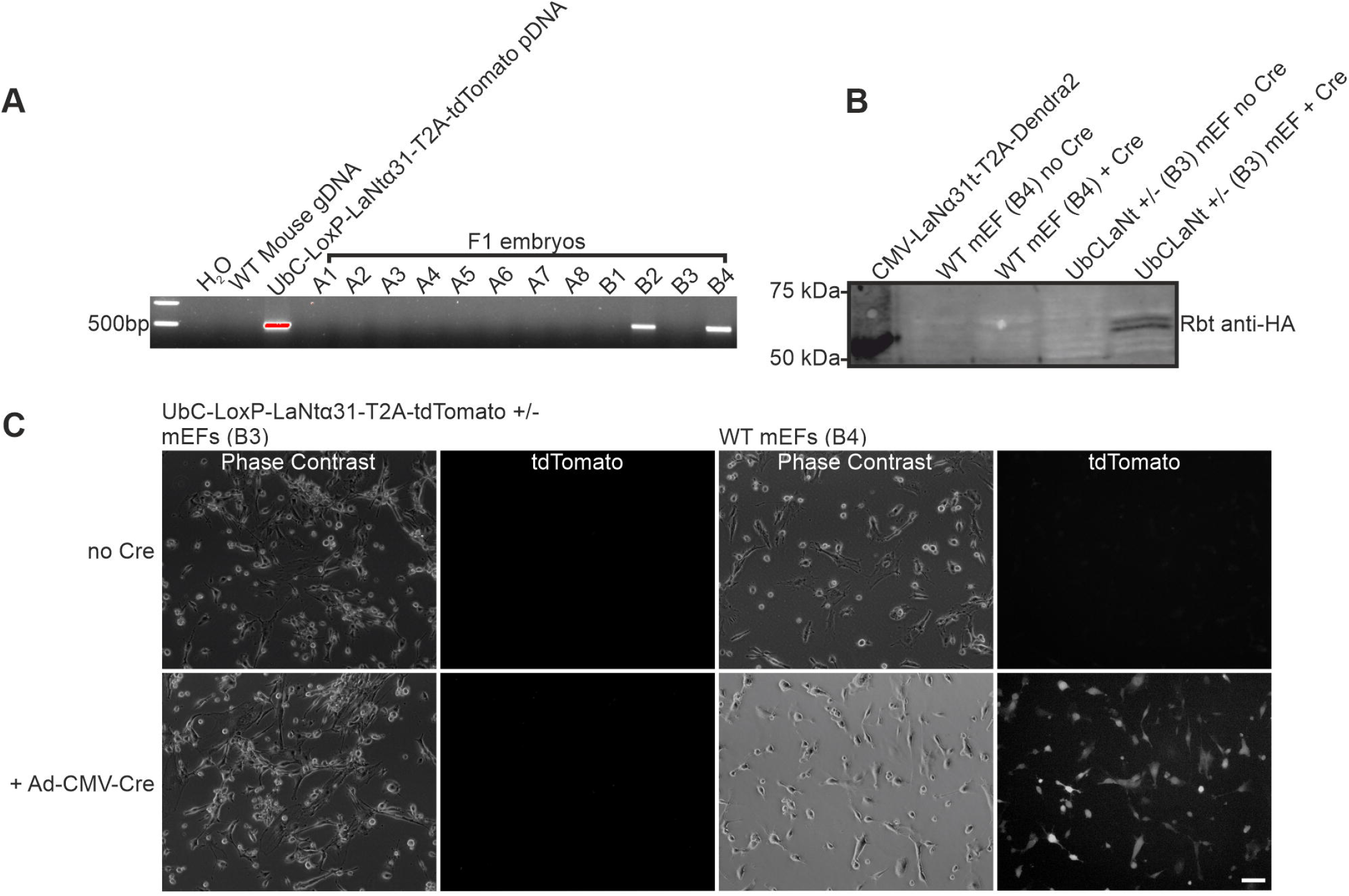
UbC-LoxP-LaNt-α31-T2A-tdTomato embryonic fibroblast express the transgene upon transduction with a Cre recombinase-coding adenovirus. A) PCR was performed on gDNA of F1 UbC-LoxP-LaNt-α31-T2A-tdTomato embryos. B) Western blot of protein lysates from explanted F1 mouse embryonic fibroblasts processed with anti-HA antibodies. C) Fluorescence microscopy images of explanted cells from UbC-LoxP-LaNt-α31-T2A-tdTomato F1 mice. Scale bar = 100 μm

Male UbCLaNt mice were mated with females from the tamoxifen-inducible ubiquitous Cre line R26CreERT2 (Fig. 3A). Transgene expression was induced by IP of tamoxifen at E13.5, and embryos collected at E19.5. PCR confirmed that Cre/LoxP mediated recombination only occurred in the embryos with both the UbCLaNt and the R26CreERT2 (Fig. 3B). Explants were generated from the skin of these embryos, and only the explants grown from double transgenic embryos exhibited tdTomato expression by fluorescence microscopy (Fig. 3C) and HA-tagged LaNt α31 expression by western immunoblotting (Fig. 3D). Together, these data confirmed the generation of tamoxifen-inducible LaNt α31 overexpressing mouse line, without detectable leakiness (UbCLaNt::R26CreERT2).

**Figure 3.**
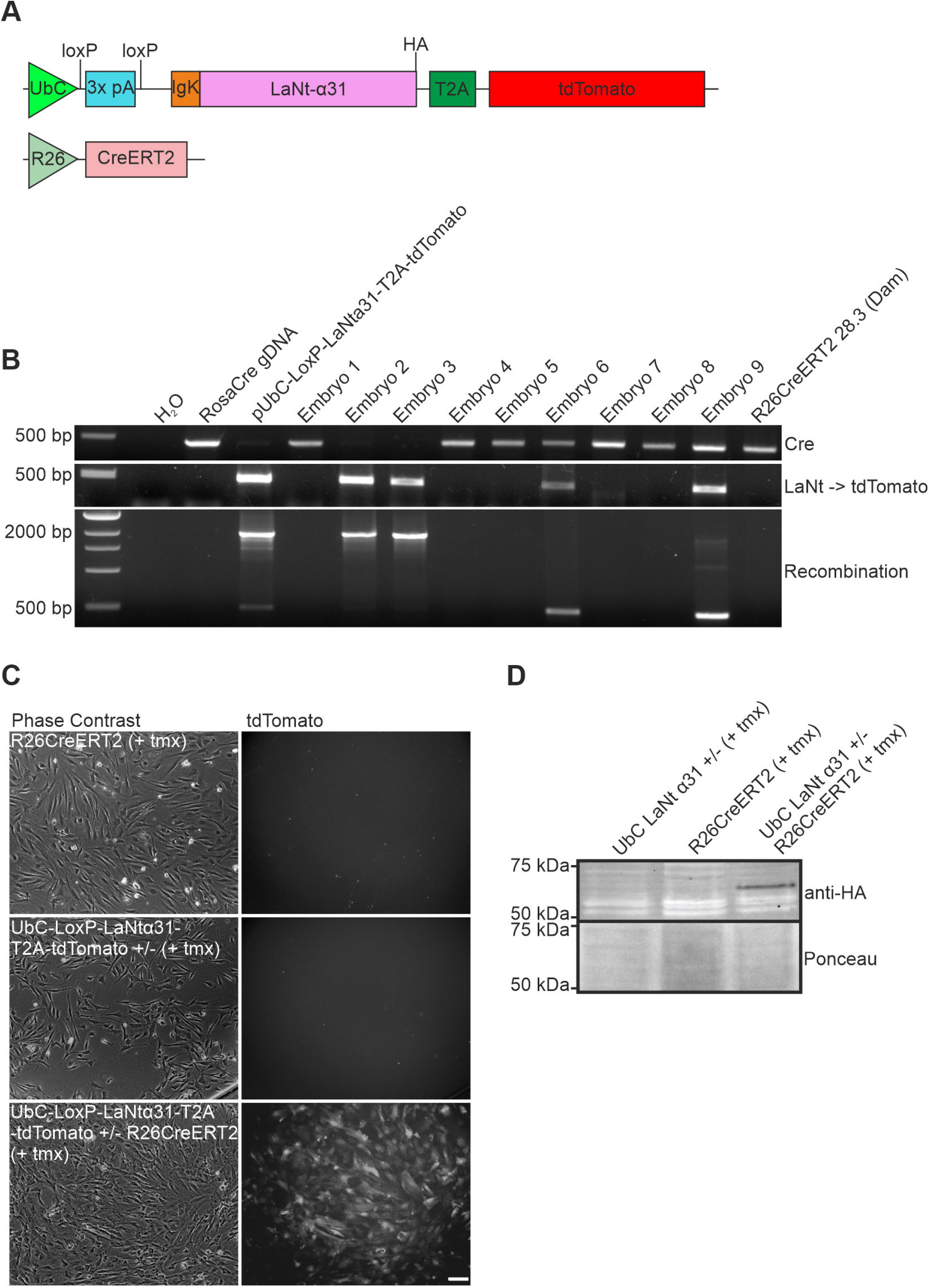
UbCLaNtα31 x R26CreERT2 ER transgenic mice express the UbC-LaNtα31 transgene following exposure to tamoxifen. A) Schematic diagram of the UbC-LaNt-α31 and Rosa-Cre transgenes. B) PCR performed using primers flanking the stop cassette on DNA extracted from transgenic mouse embryos from a UbCLaNtα31 x R26CreERT2 mating. C) Phase contrast and fluorescence microscopy images of explanted cells from UbCLaNtα31::R26CreERT2 embryos. Scale bar = 100 μm. D) Western blot of lysates from UbCLaNtα31::R26CreERT2 embryo explants processed with anti-HA antibodies.

### UbCLaNt::R26CreERT2 expression in utero causes death and localised regions of erythema at birth

To determine the impact of LaNt α31 during development where extensive BM remodelling occurs, tamoxifen was administered via IP to pregnant UbCLaNt::R26CreERT2 mice at E15.5 and pregnancies allowed to continue to term. Across two litters from different mothers, two from six pups and three from five pups respectively were intact but not viable at birth, while the remaining littermates were healthy. The non-viable pups displayed localised regions of erythema with varying severity between the mice, but were otherwise fully developed and the same size as littermates (Fig. 4A). Genotyping identified that all offspring possessed both the UbCLaNt and R26CreERT2 transgenes (Fig. 4B). Hereafter, non-viable pups are referred to as UbCLaNt::R26CreERT2 1E1, 1E2, 2E1, 2E2, 2E3, and viable pups UbCLaNt::R26CreERT2 2NE1, 2NE2. To confirm transgene expression, skin explants were established from non-viable pups, and tdTomato fluorescence was confirmed by microscopy (Fig. 4C). Consistent with the fluorescence data, western immunoblot analysis of total protein extracts from the explanted cells and whole embryo lysates revealed transgene expression in non-viable pups, although expression levels varied between the mice (Fig. 4D). To further confirm transgene expression within tissues, OCT-embedded skin sections of UbCLaNt::R26CreERT2 were processed with anti-mCherry antibodies which recognise the tdTomato protein, revealing that only the non-viable pups expressed the tdTomato reporter (Fig. 4E). Together these data confirm that only non-viable mice expressed the LaNt α31 transgene

**Figure 4.**
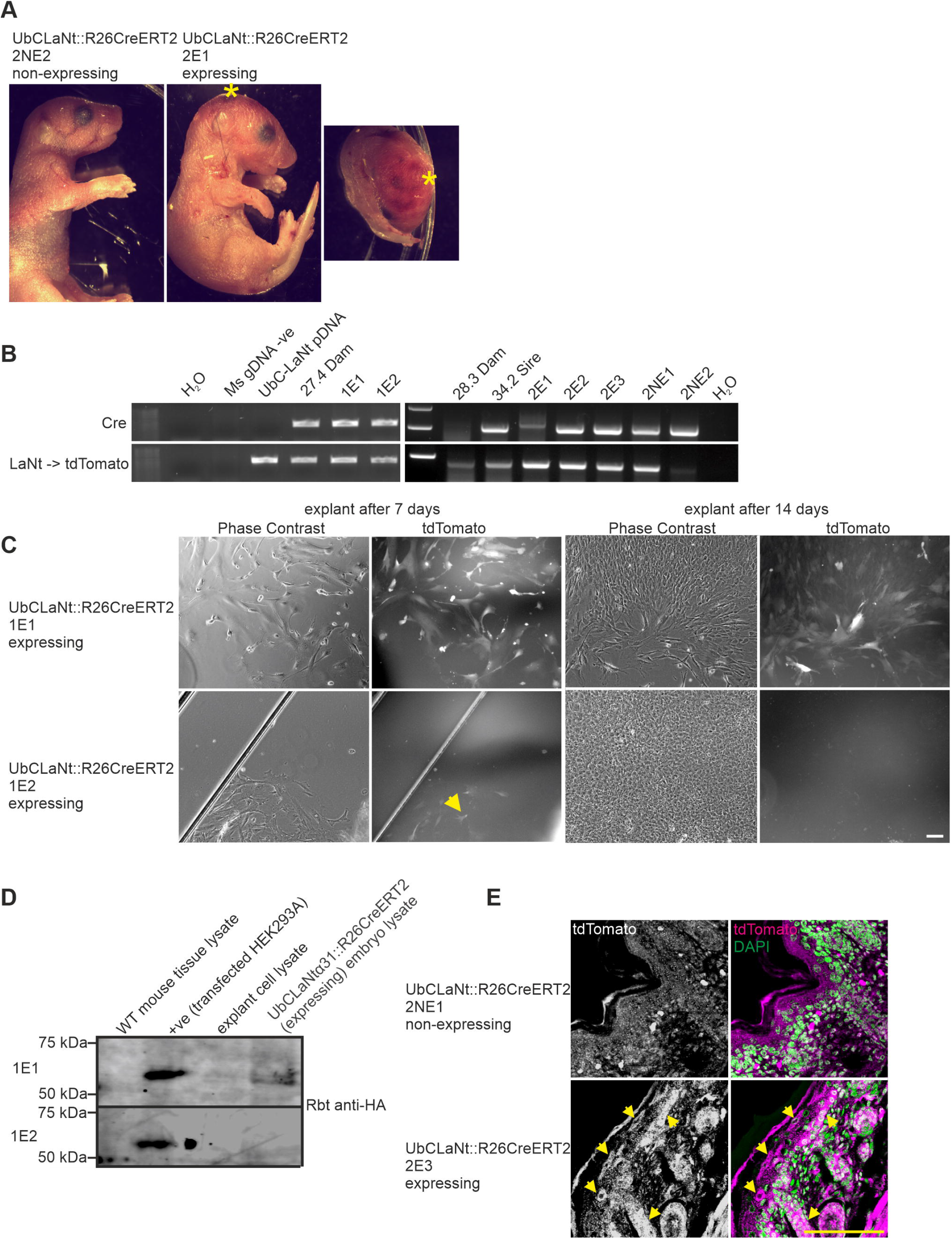
Transgenic mice overexpressing LaNtα31 display localised regions of erythema. A) Representative images of UbCLaNt::R26CreERT2 P0 mice B) PCR genotyping of transgenic mice. C) Fluorescence microscopy images of explanted cells from UbCLaNt::R26CreERT2 P0 mice. D) Western blot of tissue lysates of UbCLaNt::R26CreERT2 P0 mice. E) Representative fluorescence microscopy UbCLaNt::R26CreERT2 P0 mouse OCT sections (8 μm) probed with anti-mCherry antibodies. Yellow arrows indicate cells expressing the tdTomato transgene reporter. Scale bar = 100 μm.

To identify LaNt α31 effects at the tissue level, the pups were formalin-fixed and paraffin-embedded then processed for H&E and immunohistochemistry. All organs were present in the mice and appeared intact at the macroscopic level; however, blood exudate was observed throughout multiple tissues in all of the LaNt α31 transgene expressing mice. We focused our attention on kidney, skin and lung as examples of tissues where the BMs with differences in LM composition and where we hypothesised LaNt α31 could, therefore, elicit distinct effects. Specifically, the predominant LMs in the kidney contain three LN domains, and mutations affecting LM polymerisation lead to Pierson syndrome^19, 53–56^, whereas the major LM in the skin contains one LN domain, LM332, and loss of function leads to skin fragility, reviewed in^57^, and granulation tissue disorders^58, 59^. In the lung, LM311, a two LN domain LM, is enriched^60, 61^ and absence of LMα3 is associated with pulmonary fibrosis^62^. Each of these three tissues also express LaNt α31 in adult human tissue, and are, therefore, tissues where dysregulation of expression regulation could be physiologically relevant^38^.

### LaNt α31 overexpression leads to epithelial detachment, tubular dilation and interstitial bleeding in the kidney

In the kidneys, striking alterations were observed in the renal tubules, pelvis, and blood vessels of UbCLaNt::R26CreERT2 mice expressing the transgene. Specifically, dilation and detachment of the lining epithelia in collecting ducts and uteric bud segments was evident (Fig. 5A, black arrows), and changes were observed in the vessels of the kidney, with bleeding into the interstitial and subtubular surroundings (Fig. 5A, yellow arrows). There was some severity in the extent of the defects between the expressing pups (interstitial bleeding 4 out of 5 mice, pelvic dilation 2 out of 5, epithelial detachment and tubular observed in all mice). Indirect IF processing of tissue using antibodies raised against LM111 revealed LM localisation to be unchanged, however immunoreactivity of the tubule BMs was thickened in the expressing pups compared with littermate controls (Fig. 5B).

**Figure 5.**
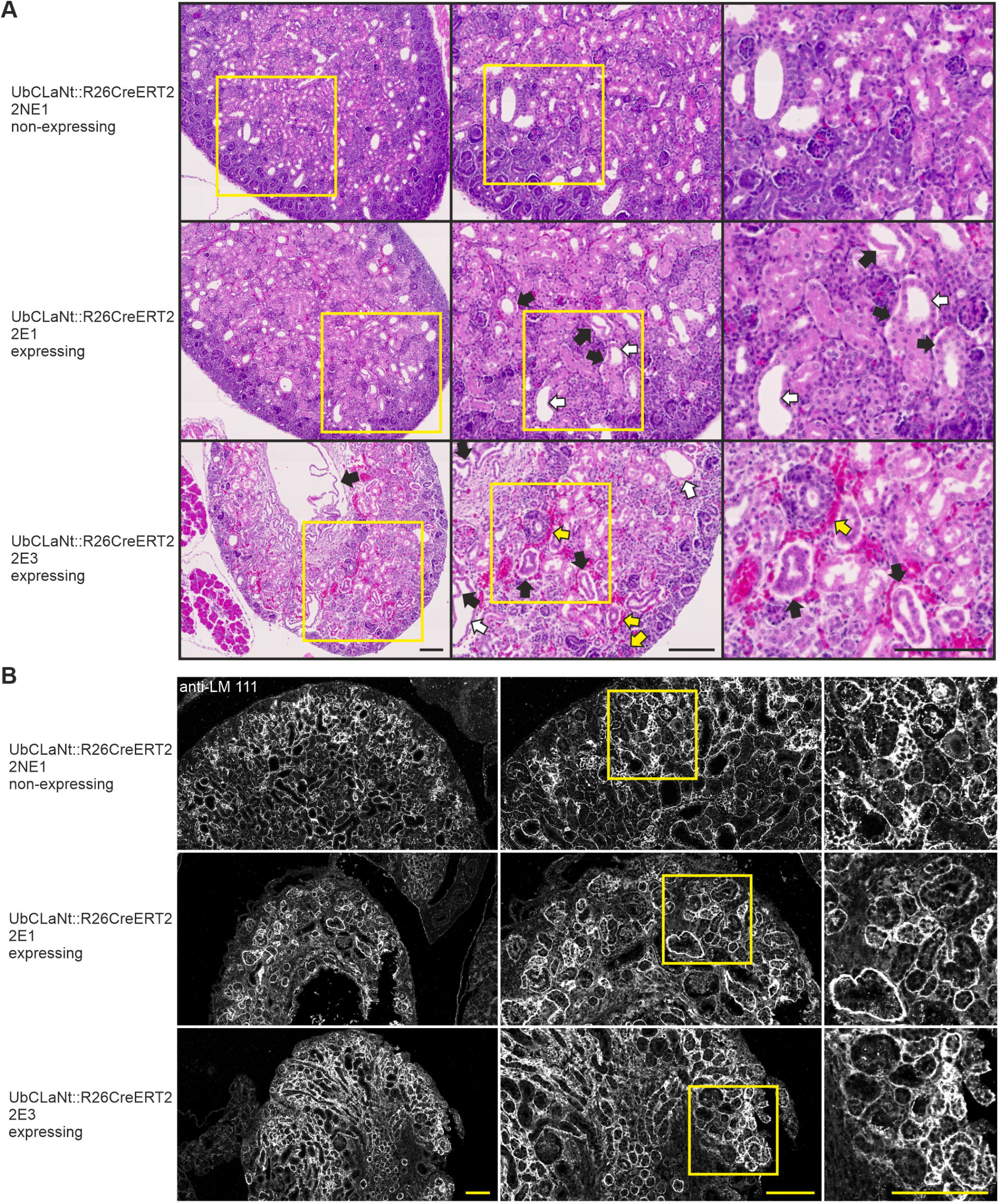
LaNt α31 overexpression leads to epithelial detachment, tubular dilation and interstitial bleeding in the kidney and thickening of the tubular basement membrane. A) Representative images of H&E stained FFPE sections (5 μm) of newborn UbCLaNt::R26CreERT2 transgenic mouse kidneys. Middle and right columns show areas of increased magnification. Black arrows point to areas of epithelial detachment. White arrows point to tubular dilation. Yellow arrows point to areas of interstitial bleeding. B) UbCLaNt::R26CreERT2 P0 mouse FFPE sections (5 μm) processed for immunohistochemistry with anti-laminin 111 polyclonal antibodies. Middle and right columns show areas of increased magnification. Scale bars = 100 μm.

### LaNt α31 overexpression disrupts epithelial basal cell layer organisation

Histological examination of the dorsal skin of UbCLaNt::R26CreERT2 mice revealed localised disruption of the epidermal basal cell layer, with a loss of the tight cuboidal structure of the stratum basale (Fig. 6A). Basal layer disruption was also observed in the outer root sheath of the hair follicles (Fig. 6A). Although no evidence of blistering at the dermal-epidermal junction was observed Mice expressing the LaNt α31 transgene displayed discontinuous LM immunoreactivity at the epidermal-dermal junction (Fig. 6B). However, the LM surrounding the outer root sheath was unaffected (Fig. 6B).

**Figure 6.**
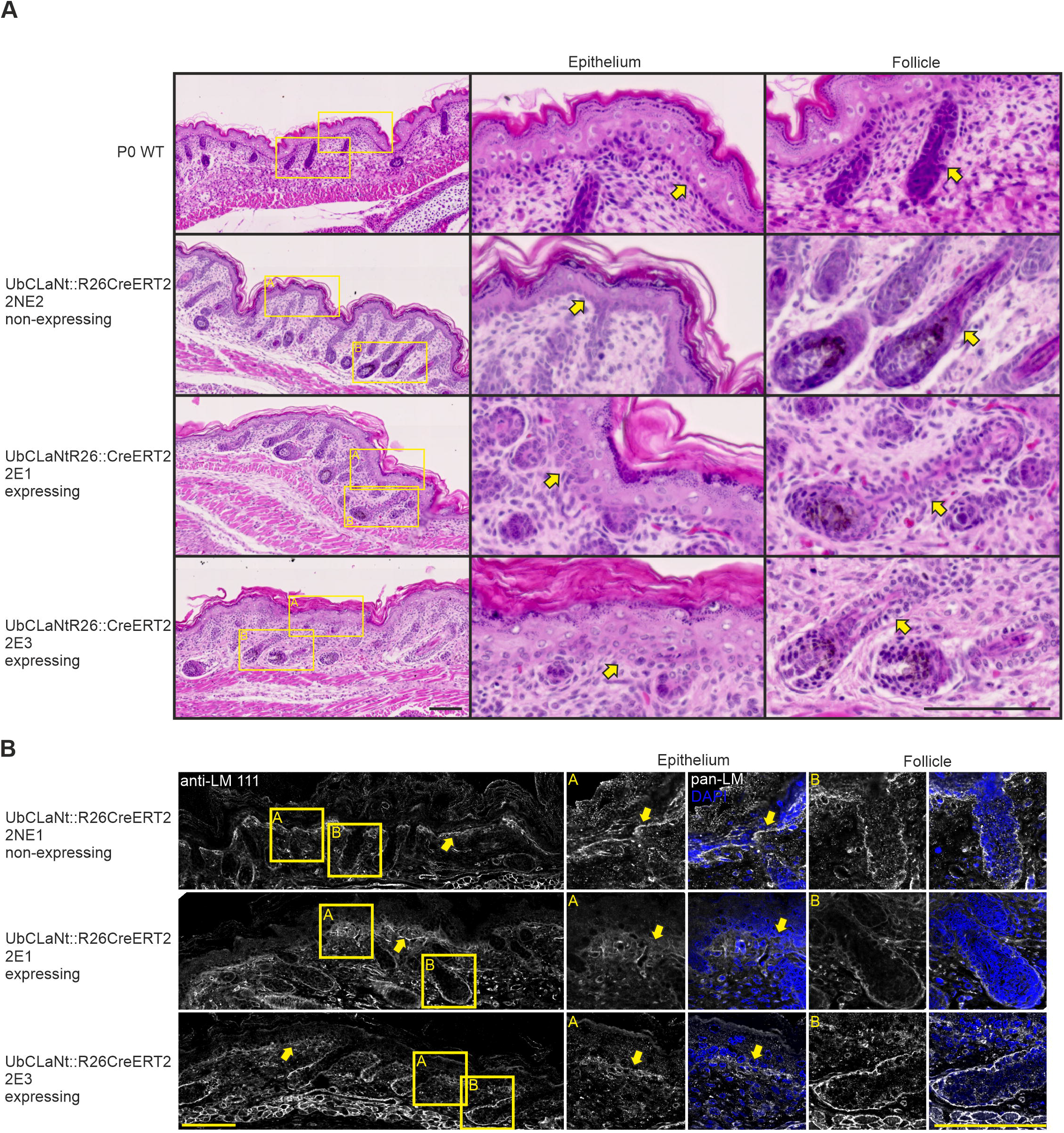
LaNt α31 overexpression disrupts epidermal-dermal basement membrane. A) H&E staining of FFPE sections (5 μm) of newborn UbCLaNt::R26CreERT2 transgenic mouse dorsal skin. Middle and right columns show increased magnification of the epithelium or hair follicles respectively. Yellow arrows indicate basal layer of epithelial cells. Scale bar = 100 μm. B) UbCLaNt::R26CreERT2 P0 mouse FFPE sections (5 μm) processed for immunohistochemistry with anti-laminin 111 immunoreactivity. Middle and right columns show areas of increased magnification. Yellow arrows indicate the epidermal-dermal junction. Scale bar = 100 μm.

### Mice expressing the LaNt α31 transgene display structural differences in the lung

Lungs of P0 mice were not inflated prior to FFPE, however structural differences between non-expressing and expressing mice were apparent. Specifically, in mice expressing LaNt α31, fewer, less densely packed alveolar epithelial cells were observed. Additionally, and similarly to the kidney, erythrocytes were present throughout the lung tissue. (Fig. 7A).

**Figure 7.**
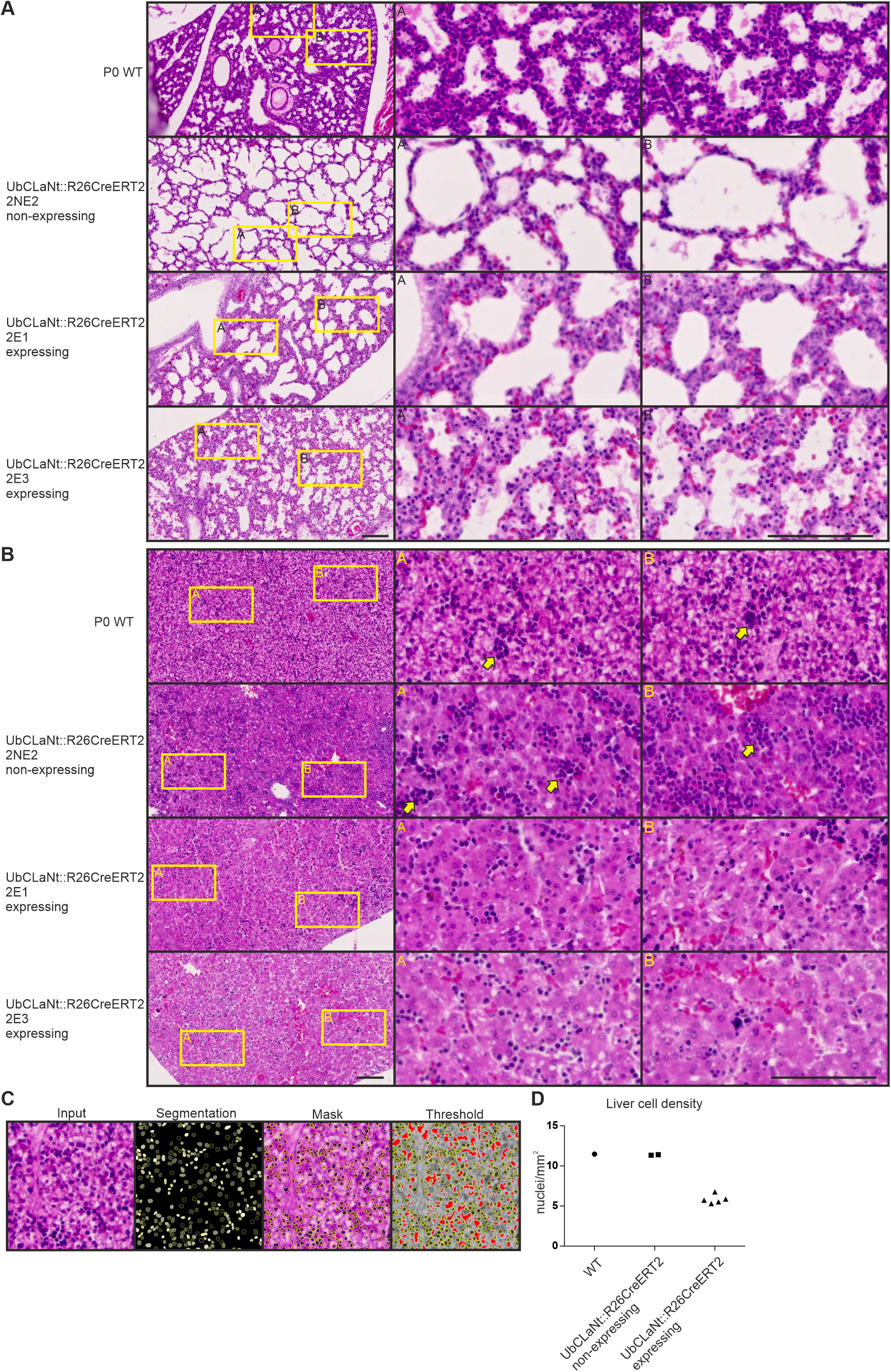
Mice expressing the LaNtα31 transgene display structural differences in the lung and a reduction of hematopoietic colonies in the liver. A) UbCLaNtα31::R26CreERT2 P0 lung FFPE sections (5 μm) stained with H&E. Middle and right columns show areas of increased magnification. B) H&E staining of FFPE sections (5 μm) of newborn UbCLaNt::R26CreERT2 transgenic mouse skin. Middle and right columns show increased magnification of different area of the liver. Yellow arrowheads highlight areas of increased cell density. Scale bars = 100 μm. C) Representative image analysis method of determining nuclei count. D) Quantification of nuclei.

### LaNt α31 overexpression leads to a reduction of hematopoietic colonies in the liver

Surprisingly, drastic and obvious superficial changes were apparent in the livers of mice expressing the LaNt α31 transgene compared to the non-expressing mice. Although the bile ducts, sinusoid endothelium and hepatocyte morphology were unchanged, there was a clear reduction in hematopoietic foci in the LaNt α31 transgene expressing animals (Fig. 7B). This reduction corresponded to a >48% reduction of total cell number (WT= 11.5 nuclei/ mm^2^, mean NE = 11.4 nuclei/ mm^2^, mean E= 5.8 nuclei/ mm^2^; Fig. 7C, D).

### Keratin 14-driven constitutive LaNt α31 induces a low offspring number

We next used a keratin-14 promoter (K14) to restrict expression to skin and the epithelia of tongue, mouth, forestomach, trachea, thymus and respiratory and urinary tracts^63–65^. K14 promoter activity has also been described in the oocyte^66^. The new construct used the human K14 promoter drive expression of human LaNt α31, followed by a T2A element and a mCherry reporter Fig. S1A) and was validated by transfecting into KERA 308 mouse epidermal keratinocytes and visualising the mCherry fluorescence (Fig. S1B) and immunoblotting for the LaNt α31 protein (Fig. S1C).

K14-LaNtα31 transgenic mice were generated by pronuclear microinjection. However, unusually small litters were obtained from recipient CD1 mothers and mice containing the transgene DNA (Fig. S1D) did not express the transgene at the protein level (Fig. S1F-G). The unusually low offspring sizes, combined with the lack of protein expression in genotype-positive mice, suggests that expression of LaNt α31 under the control of the K14 promoter is lethal during development.

## Discussion

This study has demonstrated that LaNt α31 overexpression ubiquitously during development is lethal, causing tissue specific-defects. These defects include blood exudate throughout most tissues as well as striking changes to the tubules of the kidney and the basal layer of the epidermis, depletion of hematopoietic colonies in the liver, and evidence of BM disruption at the dermal-epidermal junction. These findings build upon previous in vitro and ex vivo work that have implicated LaNt α31 in the regulation of cell adhesion, migration, and LM deposition^25,37,40^. These findings provide the first in vivo evidence that this little-studied *LAMA3*-derived splice isoform has biological significance in BM and tissue formation during development and provide a valuable platform for onward investigation.

The molecular mechanism behind the severe phenotype of the transgenic animals is challenging to determine at this point, however we can provide several plausible reasons that can explain the phenotype with substantial overlap. As LM network assembly requires binding of an α, β and γ LN domain^14–17, 67^, we predicted that the presence of an α LN domain within LaNt α31 would influence LM-LM interactions and therefore BM assembly or integrity. Consistent with this hypothesis, much of the UbCLaNt::R26CreERT2 mice phenotypes resemble those from mice where LM networks cannot form due either to LN domain mutations or overexpression of the LM-network disrupting protein, netrin-4. Specifically, mice with a mutation in the LN domain of LM α5 die before birth exhibiting defective lung development and vascular abnormalities in the kidneys^68^. While mice with LM β2 LN domain mutations or LN domain deletions exhibit renal defects, and although viable at birth, become progressively weaker and die between postnatal day 15 and 30^69–74^. Additionally, mice with netrin-4 overexpression under the control of the K14 promoter were born smaller, redder, and with increased lymphatic permeability^35^. In comparison to each of these lines, the LaNt α31 animals present with similar but more severe and more widespread phenotypes, which reflects the more widespread UBC and R26 promoter activities. Nevertheless, based on the broad similarity between these phenotypes, we propose a model where LaNt α31 overexpression inhibits LM network assembly by competing with the native LM α chain. However, within this model, there remains the question of how LaNt α31 influences tissues where there the expressed LMs do not contain an α LN domain, and therefore are not able to polymerise^16^. For example, The LM composition present within vessel BMs during development and lymph vessels is rich in the β and γ LN domain-containing LM411^75–77^. Here, one might have anticipated that the LaNt α31 LN domain could stabilise weak βγ LN dimers strengthening the BM but the observed phenotype of blood exudate throughout the mouse tissues suggests instead that the LaNt α31 transgenics have vascular leakage which overall points to a disruptive rather than stabilising role.

The in vivo findings here combined with previous in vitro studies support LaNt α31 acting as a regulator of BM homeostasis; however, we cannot yet fully rule out the possibility of LaNt α31 acting as a signalling protein^25, 37, 39, 40^. Specifically, integrin-mediated signalling from LaNt α31-like proteolytically released LN-domain containing fragments from LM α3b, α1, and β1 chains have been reported^27–29^ and some aspects of the UbCLaNt::R26CreERT2 phenotype are consistent with LaNt α31 acting in this way. For example one of the most striking phenotypes observed in the UbCLaNt::R26CreERT2 mice was the depletion of hematopoietic colonies in the liver, an essential stem cell niche during development^78–80^. Integrins α6 and β1 are highly expressed in hematopoietic stem cells, and are central to the process of migration both in and out of the fetal liver^81–83^, and a netrin-4/laminin γ1 complex has been shown to signal through the integrin α6β1 receptor^84^. Indeed, LaNt α31 may signal in a similar manner, which may be detrimental to the maintenance of hematopoietic colonies in the fetal liver. Moreover, we previously identified that LaNt α31 is enriched in human and porcine limbal stem cell niche of adult corneas, with expression further upregulated upon ex vivo stem cell activation and wound repair^37^. Coupled to the striking phenotype observed here, it is tempting to hypothesise that LaNt α31 is involved in regulating stem cell quiescence. While direct signaling effects could explain that role, indirect effects are also possible as altering LM network structural organisation changes outside-in signalling, through changing the presentation of ligands and by modifying growth factor sequestration and release rates^85^. Indeed, LM networks are critical for maintaining progenitor cell “stemness”^86–89^. Dissecting the direct versus indirect roles of LaNt α31 in intact tissue contexts is now a priority and the new transgenic line provides a valuable resource to facilitate those onward investigations.

Moving forward, the role of LaNt α31 can now be determined in a tissue and context specific manner. In this study, we focused on widespread developmental expression as a BM formation and remodelling is highly active, thereby maximising the likelihood of determining whether LaNt α31 is functional in vivo and to focus future studies upon the most relevant tissues. Considering the widespread expression of LaNt α31^38^, and the dramatic effects observed in this study, it is now important to determine effects in adult animals in normal conditions and following intervention. These studies should also include tissues where no overt LaNt α31-induced phenotype was observed. For example, although we did not observe muscle effects in these animals, LM network integrity is critical to muscle function, with the effects of LM α2 LN domain mutations or deletions developing muscular dystrophy and peripheral neuropathy with time^90–92^, therefore, longer-term studies may reveal further phenotypes once tissues are placed under stress. As the LaNt α31 phenotypes are deleterious, further studies will require lineage-specific expression to gain deeper cellular and temporal resolution.

This study provides the first in vivo evidence that LaNt α31, the newest member of the LM superfamily, is a biologically relevant matricellular protein and emphasises the importance of α LN domains as regulators of tissue homeostasis. Indeed, whereas identification of α LN domain mutations in rare inherited disorders have established that LN domains matter^19, 22, 93–95^, the LaNt α31 protein is a naturally occurring splice isoform^25^ which suggests active regulation of the LN domain interactions *via* alternative splicing. Changes to alternative splicing rates often occur in normal situations, including during development and tissue remodelling, in response to damage such as in wound repair, and can be dysregulated in pathological situations including frequently in cancer^96–98^ Considered in this way, the finding the LaNt α31 is biologically active in vivo has exciting and far-reaching implications for our understanding of BM biology.

## Abbreviations

LaNt α31: laminin N-terminus α31
BM: basement membrane
ECM: extracellular matrix
LN: laminin N-terminal
LM: laminin
LE: laminin-type epidermal growth factor-like domain
DMEM: Dulbecco’s Modified Eagle Medium
SDS-PAGE: sodium dodecyl sulfate polyacrylamide gel electrophoresis
mEFs: mouse embryonic fibroblasts
hK14: human keratin 14
intraperitoneal injection: IP

## Acknowledgements

We are grateful to the staff at the University of Liverpool Biomedical Services Unit. We would like to thank Dr. Takao Sakai, Dr. Rachel Lennon, and Dr. Mychel Morais for helpful discussions during the writing of this manuscript.

## Author contributions

**Conor J. Sugden:** Methodology, Validation, Formal analysis, Investigation, Data Curation, Writing - Original Draft, Writing - Review & Editing, Visualization. **Valentina Iorio:** Methodology, Investigation, Data Curation, Writing - Review & Editing. **Lee D. Troughton:** Methodology, Writing - Original Draft, Writing - Review & Editing. **Ke Liu:** Methodology, Writing - Review & Editing. **George Bou-Gharios:** Conceptualization, Methodology, Writing - Review & Editing, Supervision. **Kevin Hamill:** Conceptualization, Methodology, Writing - Original Draft, Writing - Review & Editing, Supervision, Funding acquisition.

---

**Supplemental Figure 1 – Transgenic expression of LaNt α31 under control of the human keratin-14 promoter results in a low number of offspring.**

A) Diagram of the phK14-LaNtα31-T2A-mCherry construct. B) Fluorescence microscopy images of KERA 308 cells transfected with phK14-LaNtα31-T2A-mCherry. C) Western blot of protein lysates from transfected KERA 308 cells. D) Schematic of F0 mice generation and PCR genotyping of F0 mice. E) PCR genotyping of F1 mice. F) Representative fluorescence images of frozen sections from F1 mice tissues. G) Western blot of tissue lysates from F1 mice, probed with anti-His antibodies.

