## Supplementary figures and images for "Laminin N-terminus α31 expression during development is lethal and causes widespread tissue-specific defects in a transgenic mouse model"

### Supplemental Figure 1

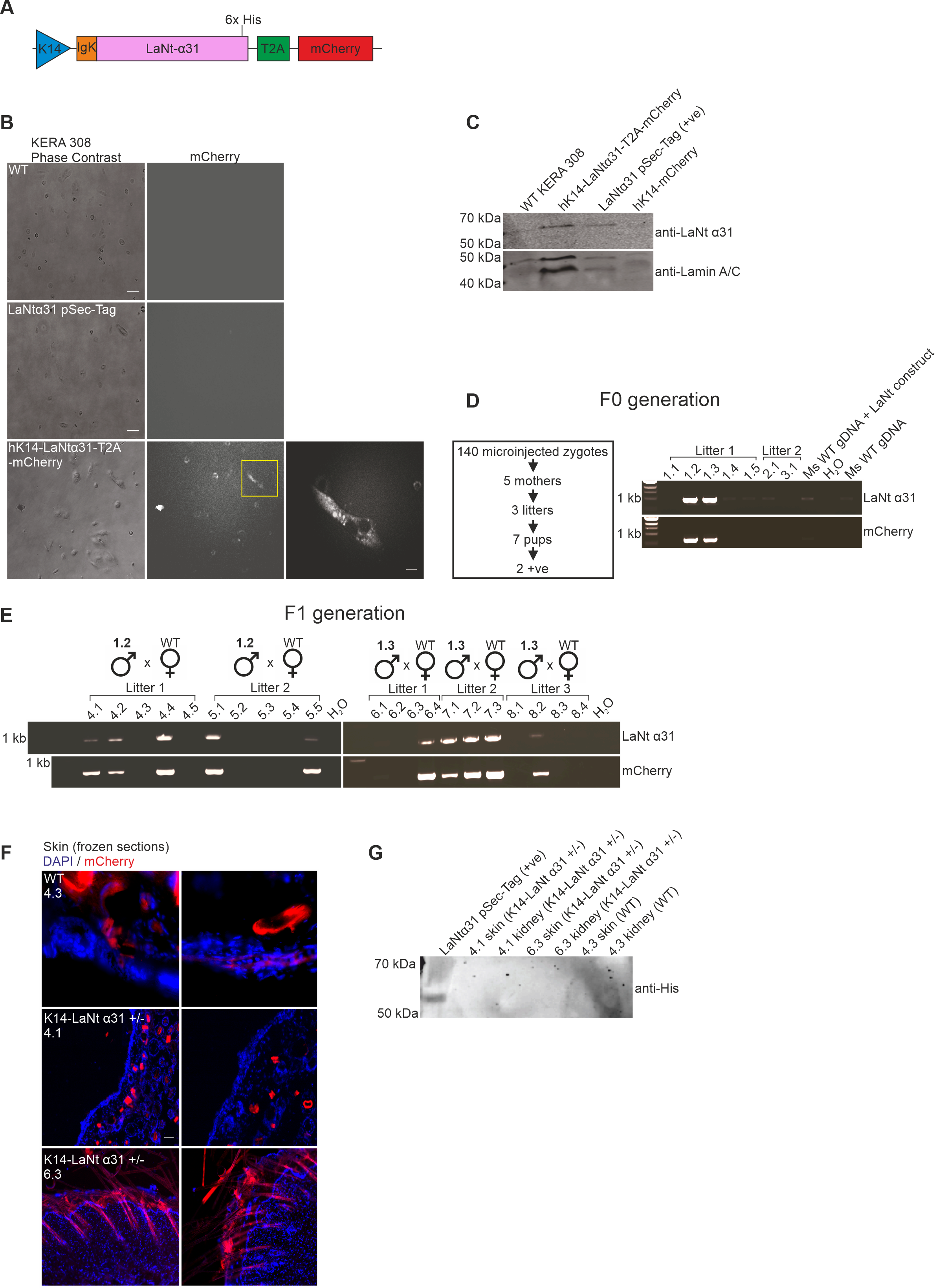
